# Temporal correlation of elevated PRMT1 gene expression with mushroom body neurogenesis during bumblebee brain development

**DOI:** 10.1101/449116

**Authors:** Cui Guan, Michaela Egertová, Clint J. Perry, Lars Chittka, Alexandar Chittka

## Abstract

Proper neural development in insects depends on the controlled proliferation and differentiation of neural precursors. In holometabolous insects, these processes must be coordinated during larval and pupal development. Recently, protein arginine methylation has come into focus as an important mechanism of controlling neural stem cell proliferation and differentiation in mammals. Whether a similar mechanism is at work in insects is unknown. We investigated this possibility by determining the expression pattern of three protein arginine methyltransferase mRNAs *(PRMT1, 4* and 5) in the developing brain of bumblebees by *in situ* hybridisation. We detected expression in neural precursors and neurons in functionally important brain areas throughout development. We found markedly higher expression of *PRMT1*, but not *PRMT4* and *PRMT5*, in regions of mushroom bodies containing dividing cells during pupal stages at the time of active neurogenesis within this brain area. At later stages of development, *PRMT1* expression levels were found to be uniform and did not correlate with actively dividing cells. Our study suggests a role for PRMT1 in regulating neural precursor divisions in the mushroom bodies of bumblebees during the period of neurogenesis.

## Introduction

The development of insects includes embryonic and postembryonic stages. The embryonic stage corresponds to the egg, whereas postembryonic stages of holometabolous insects comprise those of the larva, the pupa and the adult. During the development of holometabolous insects, the brain changes drastically not only in size but also in structure, especially during the larval and pupal stages [1]. Prominent structures of the insect brain are the optic lobes (visual system), antennal lobes (olfactory system), the central complex (which plays an essential role in sky-compass orientation [2] and aversive colour learning in honeybees [3], and in other insects regulates a wide repertoire of behaviours including locomotion, stridulation, spatial orientation and spatial memory [4, 5]), and higher order centres that coordinate sensory integration called the mushroom bodies (MBs) [6-8]. The mushroom bodies, thought to be an analogue of the mammalian hippocampus, are paired brain structures responsible for learning and memory functions in insects [9, 10]. In the adult brain, each mushroom body consists of two cap-like structures, called calyces [11], comprised of the dendrites of a large number of densely packed neurons, termed Kenyon cells [1, 11, 12]. The cell bodies of most Kenyon cells are enclosed by the calyces, while few are on the sides of or underneath the calyces [1, 11, 12].

During development of honeybees, neuroblasts located at the centre of the cups of the calyces (neuroblast clustered regions, termed proliferative regions), divide and produce Kenyon cells [1, 13, 14]. The neuroblasts begin their division from the first larval instar stage (four days from egg laying), continuing until the mid-late pupal stage (approximately five days from pupation, 16 days from egg laying) [1].

The first described mechanism for neural precursor divisions in insects involves so called Type I neuroblasts. These neuroblasts divide asymmetrically to proliferate and produce a ganglion mother cell (GMC) which undergoes a single symmetric division to produce two neurons [1]. An additional type of neuroblast (type II NBs) has been identified in *Drosophila* [15]. Type II NBs divide asymmetrically to renew themselves and generate a transit amplifying intermediate neural progenitor that continues to renew itself three to five times to generate more transit amplifying intermediate neural progenitors and a GMC that divides again to generate two neurons [15]. There are 90 type I and eight type II NBs found in each *Drosophila* brain lobe [15], where type II NBs produce many neurons for important neuropile substructures of the brain, in particular the central complex [15].

Type I neuroblast divisions occur during the development of the honeybee brain [1]. It is unclear whether type II neuroblasts are present in bees. Farris *et al*. [1] reported no evidence in honeybee mushroom bodies of neuroblast divisions other than the classic type I division pattern. However, this work was published before type II NBs were identified, leaving open the possibility that such cells might exist in bees. In this study we focus on the mushroom bodies. In *Drosophila* the neuroblasts that form the mushroom body do not undergo type II neuroblast divisions during embryonic stage, as judged by a lack of cells in this lineage that express markers of transit amplifying intermediate neural progenitors [16].

In the developing bee, neural precursor division and neurogenesis causes a dramatic change in the cytoarchitecture of mushroom bodies [1]. During the larval and pre-pupal stages the neuropils begin to form with the peduncular neuropil first observable during the third larval instar (six days from egg laying) and the calycal neuropils first seen during the pre-pupal stage (10 to 11 days from egg laying) [13]. Normally at pupal day 5, neurogenesis in the mushroom bodies of bees cease [1]. However, the growth of the mushroom body neuropil does not cease but continues throughout adult life [17]. The developmental stages of worker honeybees and bumblebees are shown in Fig. 1A, and a description of previously-reported patterns of cell division, differentiation and apoptosis during the development of mushroom bodies of honeybees is shown in Fig. 1B.

**Fig. 1.**
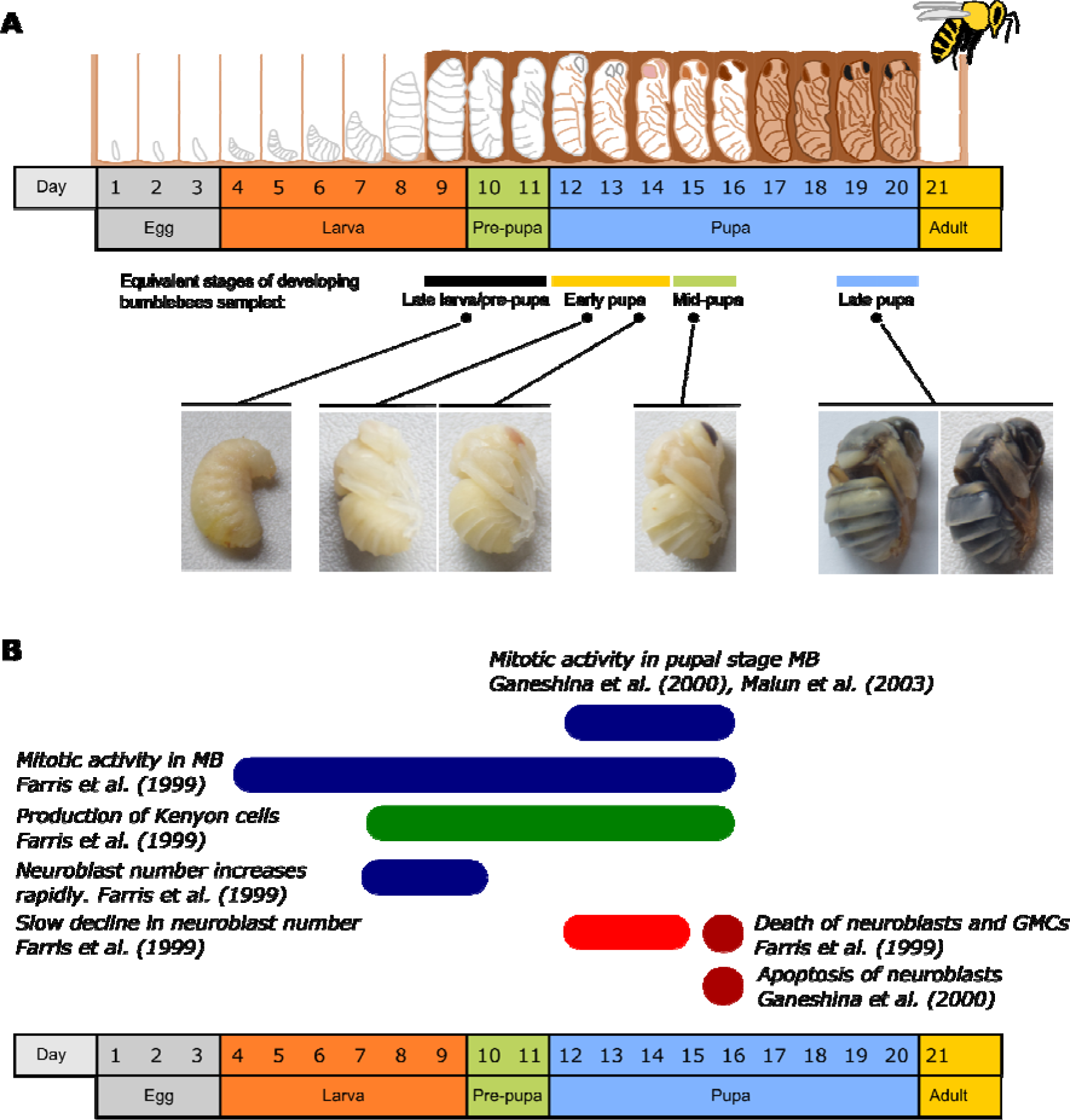
Developmental stages of honeybees and bumblebees and the corresponding changes in the composition of developing mushroom bodies (MBs). A) Schematic diagram of different developmental stages of honeybee and the equivalent stages of bumblebees investigated in this study. Given the highly close genetic and anatomic similarity between honeybees and bumblebees, eye colour and head pigmentation were used in the present study as markers to determine the equivalent bumblebee developmental stages to those stated in honeybee literature. Late larva/Pre-pupa in bumblebees correspond to days 9-11 of honeybee development; early pupa in bumblebees correspond to days 12-14 of honeybee developmental; mid-pupa in bumblebees corresponds to approximately day 15-16 honeybee developmental stage; the late bumblebee pupa corresponds to day 19-20 of honeybee development. B) Schematic diagram of reported cell division, differentiation and apoptosis during the development of mushroom bodies of honeybees. Farris *et al*. showed that there was mitotic activity in the mushroom bodies from day 4 until day 16 of development; newly produced Kenyon cells were visible from day 7 until day 16 of development; the number of mushroom body neuroblasts increases drastically from day 7 until day 9 of development, and slowly declines from day 12 to day 15 of development; cell death of neuroblasts and ganglion mother cells (GMCs) started on day 15 and was evident on day 16 of development [1]. From day 12 to day 16 of development, Ganeshina *et al*. [18] and Malun *et al*. [19] detected mitotic activity of cells in mushroom bodies. Ganeshina *et al*. showed that there were apoptotic cells from day 15 of development onwards and extensive apoptosis was observed on day 16 of development [18].

These neuroanatomical changes during bee development are well defined, but less is known about their molecular basis. In contrast, in *Drosophila*, much is known about the molecular mechanisms that control the division of neuroblasts. For example, during type I neuroblast division, the Par complex (aPKC (Atypical Protein Kinase C), Bazooka/Par3, Partitioning defective 6) forms a polarity axis in NBs [15]. The complex accumulates and segregates to the apical side. This directs the localisation of three cell fate determinants (Numb, Pros and Brat) to the opposite (basal) cortex [15]. After asymmetric division, these factors specifically segregate to the GMC, where they inhibit selfrenewal and promote differentiation [15].

Post-translational modifications of proteins play an important role in modulating their function, their interactions with various partners as well as their subcellular localisation. Different types of post-translational modifications thus fine-tune cellular responses to various environmental cues during development, allowing for stage-specific responses. Recent work has highlighted the importance of protein arginine methyltransferases (PRMTs) in regulating the development of the nervous system in vertebrates. PRMTs are a family of enzymes that catalyse the transfer of a methyl group from S-adenosylmethionine (SAM) to the guanidine nitrogen atoms of arginine [20, 21] leading to the generation of monomethyl, symmetric or asymmetric dimethyl arginines (MMA, SDMA and ADMA, respectively). ADMA is mediated by type I PRMTs, which include PRMT1, 2, 3, 4, 6 and 8, whereas SMDA is mediated by type II PRMTs, represented by PRMT5 and 9 [22, 23]. PRMTs control a multitude of essential cellular processes, e.g., cell proliferation and differentiation in all tissues during development, through the modification of protein substrates [23-30]. Their roles in controlling neural development in vertebrates are beginning to be elucidated [31-34]. For example, PRMT1 has been implicated in neurite outgrowth in human neuro2a cells during neuronal differentiation [35] and in the switch between epidermal and neural fate in *Xenopus* embryos [36]. Interestingly, enzymatic activity of PRMT1 is upregulated by the rodent antiproliferative protein TIS21 [37], a marker of all neural progenitors that are undergoing neurogenic divisions during mammalian development [38-40]. CARM1/PRMT4 regulates proliferation of PC12 cells, the cell line which is responsive to the proliferation-inducing Epidermal Growth Factor (EGF) and the differentiation-inducing Nerve Growth Factor (NGF) [41], by methylating the RNA binding protein HuD and controlling the choice of cell-cycle specific mRNAs bound by HuD in this manner [42]. PRMT5 maintains neural stem cell proliferation during early stages of development and its activity is downregulated by NGF in PC12 cells [43, 44]. Moreover, neural stem cell specific ablation of *PRMT5* in mice revealed its role in maintaining neural stem cell homeostasis during development [45].

Together, the emerging information underscores the importance of protein arginine methylation during neural development. However, very little is known about the role that PRMTs may play in insect neural development, prompting us to begin investigating their potential roles during development of the bumblebee central nervous system (CNS). Here we present the first study of the expression of *PRMT* genes during bumblebee brain development. We focus on determining the expression pattern of *PRMT1*, *PRMT4* and *PRMT5* in the brains of bumblebees at different developmental stages by *in situ* hybridisation. These genes were chosen because previous studies showed that they are important in the vertebrate nervous system development and their sequences are conserved across different species, from *Drosophila melanogaster* to mammals [22, 23, 35, 36, 42-44]. We show that all three enzymes are expressed in cell bodies in functionally important brain areas, such as the mushroom bodies, throughout development.

Our results reveal that there is a spatiotemporal correlation between levels of *PRMT1* expression and the presence of mitotically active cells in the developing mushroom body, at the time of active neurogenesis, and suggest a possible role for PRMT1 in the regulation of neuroblast/ganglion mother cell divisions.

## Materials and Methods

### Collection of bees

Bees from three *Bombus terrestris* colonies (Biobest Belgium N.V., Westerlo, Belgium) were maintained at Queen Mary University of London (Mile End campus). Pollen (approximately 7 g) and 30% sucrose (w/v) were given *ad libitum* to the hive every day during the experiment. The life cycle of a bee consists of four major stages: egg, larva, pupa, and finally the adult. Generally speaking, for a *Bombus terrestris* worker bee, eggs hatch into larvae after 4-6 days [46]. The larval stage lasts for 10-20 days before it pupates. Then the larva moults and spins a silken cocoon around its body. After about two weeks as a pupa, an adult worker emerges [46]. Six developmental stages of bees were collected. These were late larvae/pre-pupae (with the larva ceasing any movements and the cocoon being formed), early-pupae (white/pink eye pupae), mid-pupae (brown eye pupae), late pupae (the cocoon contains a black body and head), two-day old workers, and seven to ten-day old workers. Given the highly close genetic and anatomic similarity between honeybees and bumblebees [47, 48], eye colour and head pigmentation were used in the present study as markers to sample the pupal staged bees based on the honeybee literature (Fig. 1A) [49]. Two-day old workers were sampled because this is the earliest age a worker bee can start to forage [50]. Furthermore, two-day old worker honeybees without flight experience go through a drastic outgrowth of Kenyon cell dendrites in mushroom bodies [17]. Seven to ten-day old workers were sampled because this range was the average age for a bumblebee to start to forage according to the present experimental observations. In honeybees, foraging bees have more dendritic spines in the mushroom body in comparison to nursing bees, which are normally younger than foraging bees [17]. All the bees were kept inside the nest without any flight experience, to remove the possibility of flight experience causing changes in the brain of the bees that were sampled.

All bees were gently removed from the nest using large tweezers and then placed over ice to anaesthetize them before dissection. The entire heads of late larvae/pre-pupae and early-pupae were removed and placed into 4% paraformaldehyde (PFA) in phosphate-buffered saline (PBS: 0.2562g NaH_2_PO_4_·H_2_O, 1.495g Na_2_HPO_4_·2H_2_O, 8.766g NaCl per liter, pH 7.2-7.4) for fixation at 4 °C overnight. For mid-pupae, late pupae and adults, the heads were removed and the brains were immediately dissected out from the head capsule under cold 4% PFA in PBS. Subsequently, the brains of mid-pupae, late pupae and adults were put in fixative 4% PFA in PBS at 4 °C overnight. The next day the tissue was washed in PBS three times (10 min each) before being transferred to 10% sucrose/PBS and 20% sucrose/PBS for 4 hours each at room temperature and 30% sucrose/PBS overnight at 4 °C for cryoprotection. On the next day, the tissue was embedded in optimal cutting temperature compound (O.C.T; Agar Scientific Ltd, UK) and rapidly frozen on dry ice before being sectioned into 10 μm slices. Serial brain sections were alternately mounted onto positively charged SuperFrost Ultra Plus slides for later use (Fisher Scientific UK Ltd).

### Probe synthesis

*In situ* hybridisation of bumblebee brain cryosections was conducted with digoxigenin (DIG)-labeled riboprobes. For *PRMT1*, antisense and sense probes were transcribed from a pBluescript II SK (+/−) subclone containing a 575-bp fragment from the coding DNA sequence (CDS) region (bp 539-1113; NCBI RefSeq: XM_003395460.2) of the *PRMT1* cDNA. DNA containing the *PRMT1* fragment flanked by T3 RNA polymerase and T7 RNA polymerase sites was amplified by PCR. The antisense probe was synthesized using T3 RNA polymerase, whereas the sense probe was synthesized using T7 RNA polymerase. For *PRMT4/CARMER*, antisense and sense probes were transcribed from a pcDNA3 subclone containing a 517-bp fragment from the 3’-untranslated region (bp 2364-2880; NCBI RefSeq: XM_012313193.1) of the *PRMT4* cDNA. DNA containing the *PRMT4* fragment flanked by SP6 RNA polymerase and T7 RNA polymerase sites was amplified by PCR. The antisense probe was synthesized using SP6 RNA polymerase, whereas the sense probe was synthesized using T7 RNA polymerase. For *PRMT5*, antisense and sense probes were synthesized from a 463-bp fragment from the CDS region (bp 1626-2088; NCBI RefSeq: XM_003396560.2) of the *PRMT5* cDNA pcDNA3 clone. DNA containing the *PRMT5* fragment flanked by SP6 RNA polymerase and T7 RNA polymerase sites was amplified by PCR. The antisense probe was synthesized using SP6 RNA polymerase with a PCR amplified fragment from the plasmid, whereas the sense probe was synthesized using T7 RNA polymerase. A DIG RNA Labelling Kit (Roche, UK) was used to synthesize the DIG-labeled riboprobes according to the manufacturer’s instructions.

### *In situ* hybridisation (ISH)

Frozen horizontal brain sections were air dried for two hours and then washed twice for 7 minutes in PBS before being post-fixed in 4% PFA at room temperature for 20 minutes. This was followed by three 5-minute washes with PBS containing 0.1% Tween20 (Sigma-Aldrich, UK) (PBST). Slides were then incubated at 37°C for 7 minutes in 20 mg/mL Proteinase K (Roche, UK) in Tris/EDTA (6.25 mM EDTA, 50 mM Tris, pH 7.5) to increase probe penetration and then were put into 4% PFA for 5 minutes to prevent the sections from falling apart. The sections were then washed in PBST three times (5 minutes per time), and acetylated in an acetylate solution (0.1M Triethanolamine (TEA), 0.25% acetic anhydride, 0.175% acetic acid) for 10 minutes in order to remove the charge on the sections and eliminate the background binding of the probes later. This was followed by two 5-minute PBST washes and one 5-minute wash in 5* saline sodium citrate (SCC; Na-citrate 0.075 M, NaCl 0.75 M, pH 7). Sections were then incubated in pre-hybridisation buffer (50 μg/mL yeast RNA, 50% Formamide, 20% 20*SSC, 50 μg/mL Heparin and 0.1% Tween 20) for two hours at room temperature. Subsequently, hybridisation was performed by incubating the sections overnight at 65 °C with hybridisation buffer, which contained a DIG-labeled ribo-probe of either antisense or sense of *PRMT1, 4* and *5* at a concentration of 1 μg/ml. On the following day, slides were equilibrated in 5* SSC once for 20 minutes and 0.2* SSC for 40 minutes twice at 65 °C. This was followed by a 10 minute 0.2* SSC wash and a 10-minute buffer B1 (5M NaCl, 1M Tris, pH 7.5) wash at room temperature. Slides were then blocked in buffer B1 with 5% goat serum for two hours at room temperature, and incubated with Anti-Digoxigenin-AP Fab Fragments (Roche, UK) antibody diluted 1:3000 in buffer B1 with 2.5% goat serum overnight at 4°C. On the next day, the slides were washed with buffer B1 three times and once with buffer B3 (1M MgCl_2_, 5M NaCl and 1M Tris, pH 9.5). Products were visualized using NBT/BCIP reagent (Roche, UK) according to the manufacturer’s instructions in buffer B3 with 0.1% of Tween 20 (Sigma-Aldrich, UK) in a dark and humidified chamber. The staining was detected by the presence of dark purple precipitate. Conditions for colour development were kept identical for all experiments performed with each sense and antisense probe. Colour development was monitored by using a LEICA DMR4 microscope (LEICA, Germany) every half an hour to determine when to stop the reaction. Reactions were stopped by putting the slides in deionised water when a moderate intensity of staining was achieved.

Slides were then mounted. For the mounting procedure, the slides were washed with deionised water three times (5 minutes each), and subsequently dried for about 45 minutes at 37 °C until they were totally dry before being put in 100% ethanol to dehydrate twice for 10 seconds, and equilibrated in histo-clear (National Diagnostics, UK) twice (7 minutes each). Histomount (National Diagnostics, UK) was then added to the slides before cover slips were added. Following this, the slides were allowed to dry and stored in a dark box.

Sections were photographed with a QIMAGING QIClick™ CCD Colour Camera linked to a DMRA2 light microscope (LEICA, Germany) using image analysis software (Volocity^®^ software, v.6.3.1, PerkinElmer, USA) running on an iMac (27-inch, Version 10.10, Late 2013 model with OS X Yosemite). Photographs of half brain sections were collected (the other half of the brain showed symmetrical staining). Adjacent sections from the same brain were used for sense and antisense probes to ensure the specificity of the staining with antisense probes.

### Quantification of ISH data for *PRMT1*

To quantify relative intensities of mRNA expression, ImageJ was used to determine the optical density of selected regions as a measure of gene expression [51]. Thirty areas from the mushroom bodies, optic lobes, antennal lobes and background (no *PRMT1* ISH staining) neuropil areas were chosen for analysis. The goal was to choose areas that were from equivalent anatomical areas across the different sections and time-points. In figure 5A and B, areas 1-4 are the central mushroom bodies, areas 5-12 are the peripheral mushroom body, areas 13-18 are areas alongside the antennal lobe, and areas 19-24 are within the optic lobe area. In addition, a high intensity sub-region of the antennal lobe (area 13) was selected because in the early pupal stage this area repeatedly showed darker staining (we termed this the ‘antennal lobe high intensity region’). Areas 25-30, within the neuropil region that lacked detectable staining, were chosen as the background areas. An example of the areas chosen are shown in Fig. 5A (original image of a bee section) and B (image showing the selected regions used for quantification). Areas high in noise (such as a folded section area or non-removable dirt on the cover slip) were avoided when choosing the areas of interest.

ImageJ was used to measure the optical density of the region of interest (ROI) [51]. The average optical density of the background areas was subtracted from each ROI to normalize the intensity of the staining in each section. Five grouped areas (mushroom body central areas, mushroom body peripheral areas, antennal lobe high intensity area in early pupal stage, antennal lobe area and optic lobe area) were compared (see, for example, Fig. 5B).

### Immunohistochemistry

Tissue sections which were hybridized with the *PRMT1* specific antisense riboprobe were then used for immunohistochemistry. The slides were washed with PBS and blocked in 5% bovine serum albumin (BSA; Sigma-Aldrich, UK) /PBS-0.1% Triton X-100 (Sigma-Aldrich, UK) (PBS-Tx containing 5% BSA) at room temperature for 1 hour, and incubated with 5% BSA/PBS-Tx containing 1:500-fold diluted anti-Phospho-Histone H3 [pSer10] antibody (PH3S10; produced in rabbit; Sigma-Aldrich, UK) at 4 °C overnight. The next day the slides were washed three times in PBS (10 minutes for each wash) and incubated with a Cy2-conjugated anti-rabbit IgG secondary antibody at 1:500 dilution (Jackson Immunoresearch, UK) and Hoechst 33258 (1 μg/ml; Sigma-Aldrich, UK) for 2 h. This was followed by three washes with PBS-Tx (10 minutes each). Images were analysed using a fluorescence microscope (LEICA DMRA2, LEICA, Germany) and photographed with a digital camera (HAMAMATSU, ORCA-ER, C 4742-80, HAMAMATSU, Japan). The photographs were saved in Volocity software (v.6.3.1, PerkinElmer, USA) running on an iMac (27-inch, Version 10.10, Late 2013 model with OS X Yosemite, Apple, USA).

### Correlation analysis of *PRMT1* expression with mitotically-active cells and cell density within mushroom bodies

The optical density of staining associated *PRMT1* mRNA expression was compared with the proportion of mitotically active cells measured as the number of anti-PH3S10 positive nuclei divided by the total number of nuclei (identified by Hoechst staining) in each selected region of the mushroom bodies.

To do this, the ISH image was used to select five ROIs in the dark stained region (central region) of the mushroom body and five ROIs in the left and right-side regions (peripheral regions) of the mushroom body to analyse. Two regions of background in the non-stained neuropil were also chosen to normalise the optical density of ISH image. The ISH channel was used to select these regions to avoid any bias involved in selecting the mitotic marker stained regions. The optical density of each selected region and the region size were measured using Fiji ImageJ software [51]. The corresponding numbers of nuclei stained with Hoechst and numbers of PH3S 10-expressing cells in the same selected regions were counted. An example of how ROIs were selected is shown in Supplemental Fig. S1.

Spearman’s rank correlation coefficient analysis was used to analyse the correlation between the expression level of *PRMT1* (measured as optical density normalised by the subtraction of the background optical density) and proportion of mitotically active cells (the number of PH3S10-positive dividing nuclei divided by the total number of nuclei) in each selected region for three developmental stages of bumblebees (late larvae/pre-pupae, early-pupae and mid-pupae).

Furthermore, the Hoechst stained image and the ISH image was analysed to gain a better understanding of the relationship between the cell density (*measured as the number of nuclei divided by the size of the area*) and the expression level of *PRMT1* in the same region. Spearman’s rank correlation coefficient analysis was also used to determine whether there is a spatio-temporal correlation between *PRMT1* mRNA expression level and the cell density during the three stages of bumblebee development. R statistical software version 3.4.0 was used in the analyses above [52].

## Results

### *In situ* hybridisation analysis of *PRMT* gene expression in pupal and adult bumblebee brains

To gain insights into potential roles of PRMTs in bumblebee neural development, we performed *in situ* hybridisation to determine the mRNA expression pattern of *PRMT1*, *PRMT4* and *PRMT5* in the developing brain of bumblebees. We selected the late larval/pre-pupal, early-pupal, mid-pupal, late pupal, two-day old worker, and 7 to 10-day old worker stages to investigate mRNA expression of *PRMT1*, *4* and *5*, since the first three developmental stages are marked by active neurogenesis in the mushroom body, while it ceases after the mid-pupal stage [1].

Our results show that all three genes are expressed throughout pupal development and adult stages in cell bodies of neural precursors and nerve cells in the mushroom bodies, antennal lobes, and optic lobes (Fig. 2, 3 and 4). Interestingly, we observed that the expression of all *PRMTs* was much more restricted in the adult brain (Fig. 2, 3 and 4 A, B, and C) than their broader expression in the pupal brain (Fig. 2, 3 and 4 D, E, and F), suggesting a role for these enzymes in pupal brain development. Examples of images of sections which were hybridized with sense controls are shown in the inset panels of Fig. 2F, 3F and 4F, confirming the specificity of our ISH probes.

**Fig. 2.**
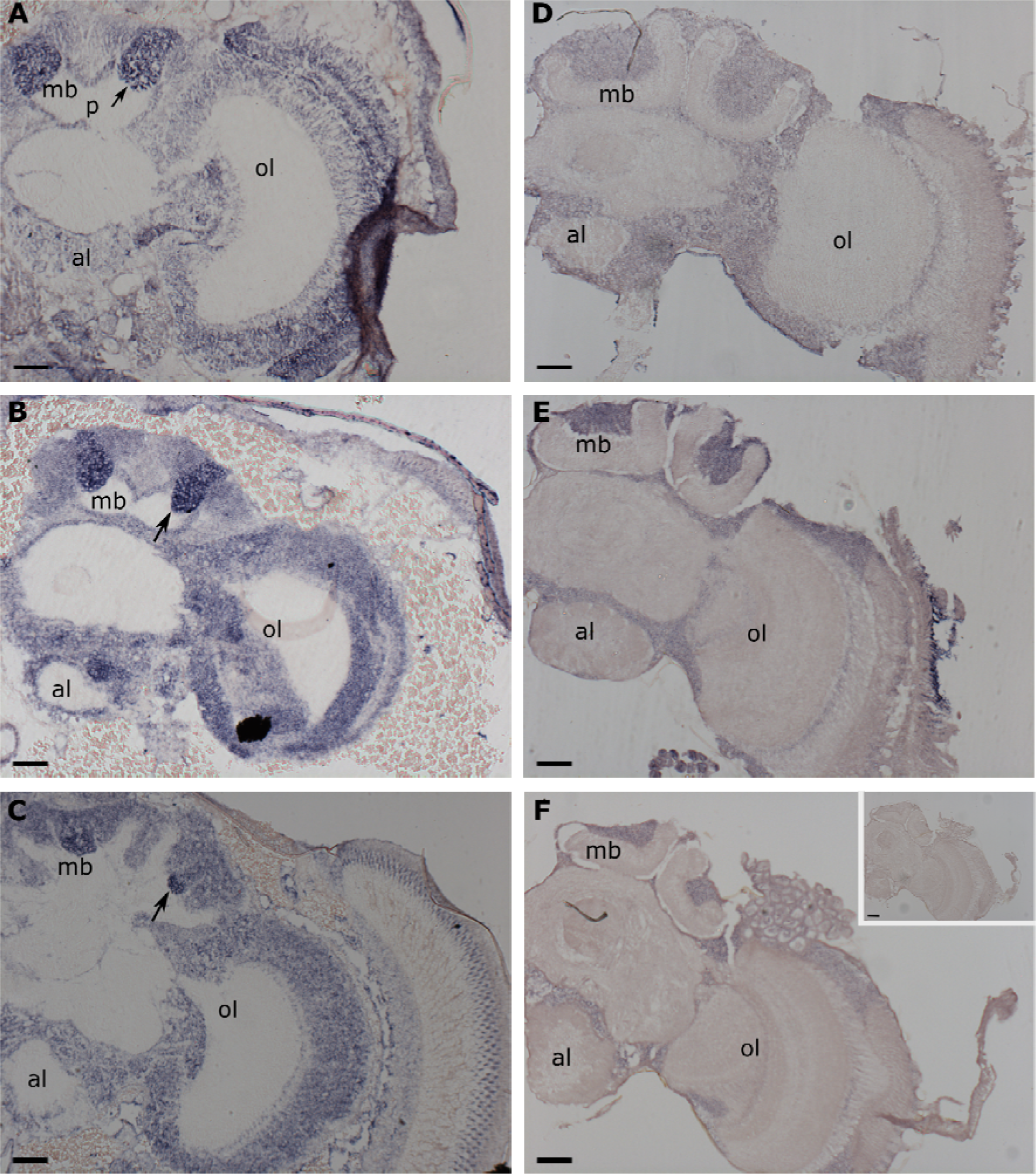
*In situ* hybridisation of *PRMT1* in the frontal half section of the brain of A) late larva/pre-pupa; B) early pupa; C) mid-pupa; D) late pupa; E) two-day old worker and F) 7 to 10-day old worker. Three biological replicates were analysed for each stage. Arrow: neural precursor dividing regions according to honeybee literature. mb: mushroom body, ol: optic lobe; al: antennal lobe; p: pedunculus. F) inset panel shows sense control without signal. Scale bars: 120 μm.

**Fig. 3.**
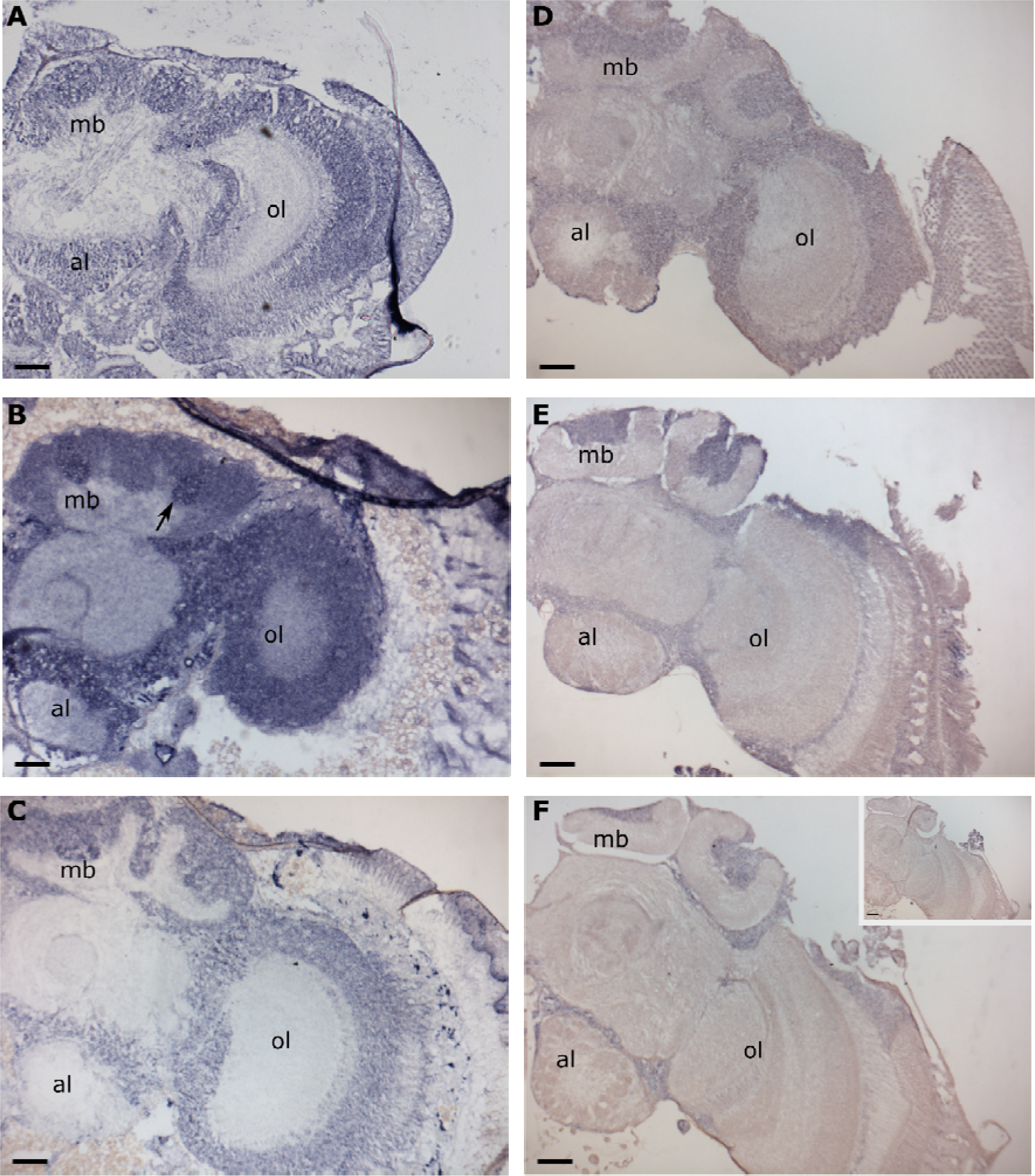
*In situ* hybridisation of *PRMT4* in the frontal half section of the brain of A) late larva/pre-pupa; B) early pupa; C) mid-pupa; D) late pupa; E) two-day old worker and F) 7 to 10-day old worker. Three biological replicates were analysed for each stage. Arrow: higher expression region. mb: mushroom body; ol: optic lobe; al: antennal lobe. F) inset panel shows sense control without signal. Scale bars: 120 μm.

**Fig. 4.**
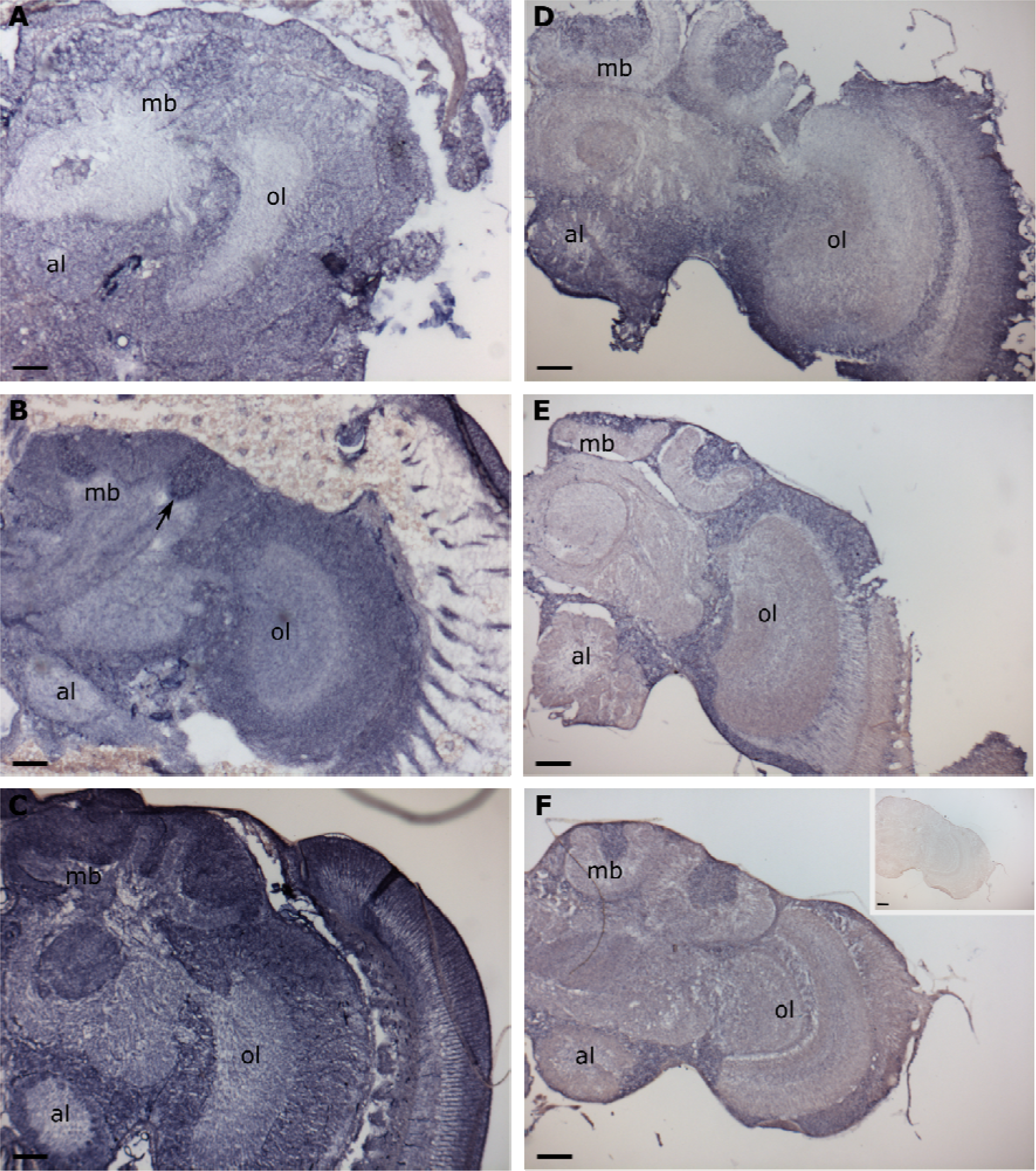
*In situ* hybridisation of *PRMT5* in the frontal half section of the brain of A) late larva/pre-pupa; B) early pupa; C) mid-pupa; D) late pupa; E) two-day old worker and F) 7 to 10-day old worker. Three biological replicates were analysed for each stage. Arrow: higher expression region. mb: mushroom body; ol: optic lobe; al: antennal lobe. F) inset panel shows sense control without signal. Scale bars: 120 μm.

#### PRMT1 mRNA expression

The expression level of *PRMT1* mRNA at the developmental stages of late larvae/pre-pupae, early-pupae and mid-pupae was higher in the mushroom body central regions, which are thought to contain dividing neural precursors (arrows in Fig. 2), than in the mushroom body peripheral regions, which are thought to contain differentiated neurons, at least in honeybees (Fig. 2A, B and C) [1, 13]. Previous studies in honeybees have shown that neuroblast clusters localize at the central region of mushroom bodies where they divide before they differentiate and migrate to the periphery of the neuroblast clusters [1, 13]. The newly born post-mitotic neurons, called Kenyon cells, are located at the periphery of the neuroblast clusters [1, 13]. The stronger localized expression of *PRMT1* mRNA in central mushroom body areas during earlier stages of development contrasted with its uniform expression pattern in the late pupal and adult stages (Fig. 2D, E and F).

#### PRMT4 and PRMT5 mRNA expression

In contrast to *PRMT1*, *PRMT4* and *PRMT5* mRNAs were more uniformly expressed across all stages (Fig. 3 and 4) except for the early pupal stage (Fig. 3B and Fig.4B). There was a slightly higher expression level of *PRMT4* and *PRMT5* in central mushroom bodies at early pupal stage (Fig. 3B and Fig. 4B), although this was less distinct than the markedly elevated level of expression observed for *PRMT1*. No significant differences in expression levels of either *PRMT4* or *5* were detected in any of the sub-regions of the mushroom bodies, optic lobes or antennal lobes at other developmental stages investigated (*PRMT4*: Fig. 3A and C-F, *PRMT5*: Fig. 4A and C-F).

### Quantification of *PRMT1* expression as revealed by mRNA *in situ* hybridisation

We noticed that *PRMT1* mRNA was preferentially expressed in the central mushroom body regions in pre-, early and mid-pupal stages (Fig. 2A, B and C), at the time when neurogenesis occurs in the honeybee [1]. Given the importance of central mushroom bodies for neuroblast/ganglion mother cell divisions we sought to quantify the relative expression level of *PRMT1* mRNA within different selected regions of the CNS. To this end, we measured the optical density of 24 areas of the CNS grouped into 4 anatomical regions (Fig. 5A and B and see also Materials and Methods). For the early pupal stage an additional high intensity sub-region of the antennal lobe was also chosen and analysed. The results are shown in Fig 5C. *PRMT1* expression was higher in the central mushroom body areas and a sub area of the antennal lobe than in other areas during the late larvae/pre-pupae, early pupae and mid-pupae developmental stages. The higher *PRMT1* mRNA expression level within these areas of mushroom bodies and antennal lobe during these developmental stages suggests that there may be a functional significance to this higher level of expression.

### High levels of *PRMT1* expression are found in areas of mitotically active cells within the mushroom bodies

*PRMT1* was preferentially expressed in the centre of the mushroom bodies, the area which is recognised as a region containing dividing neuroblasts and ganglion mother cells from larval to midpupal stages in honeybees [1, 13, 14]. Therefore, we investigated whether the same anatomical areas also contain dividing cells in bumblebees. To this end, we used an antibody that marks mitotically active cells to co-immunolabel sections of the bumblebee brains used for ISH analysis of *PRMT1* mRNA expression to determine if higher levels of *PRMT1* mRNA correlate with the prevalence of mitotically active cells. The antibody against phosphorylated serine 10 on histone H3 (PH3S10) is specific for the metaphase and anaphase stages of mitotically active cells [53]. We found a significant number of cells within the mushroom bodies of the late larva/pre-pupa, early pupa, and mid-pupa regions, shown by ISH to have higher expression of *PRMT1*, to be mitotically active (Fig. 6M-O). The observed number of mitotically active cells is likely to underestimate the total number of dividing cells because PH3S10 only marks the metaphase and anaphase stages of the cell cycle [53].

In contrast, in late pupal and adult stages investigated here, there was a more uniform expression pattern of *PRMT1*, and we did not detect any cells expressing PH3S10 (Supplementary Fig. S2). These observations suggest that there were no dividing cells in the adult brain, consistent with previous observations that demonstrated the absence of adult neurogenesis in honeybees [1, 13, 54].

**Fig. 5.**
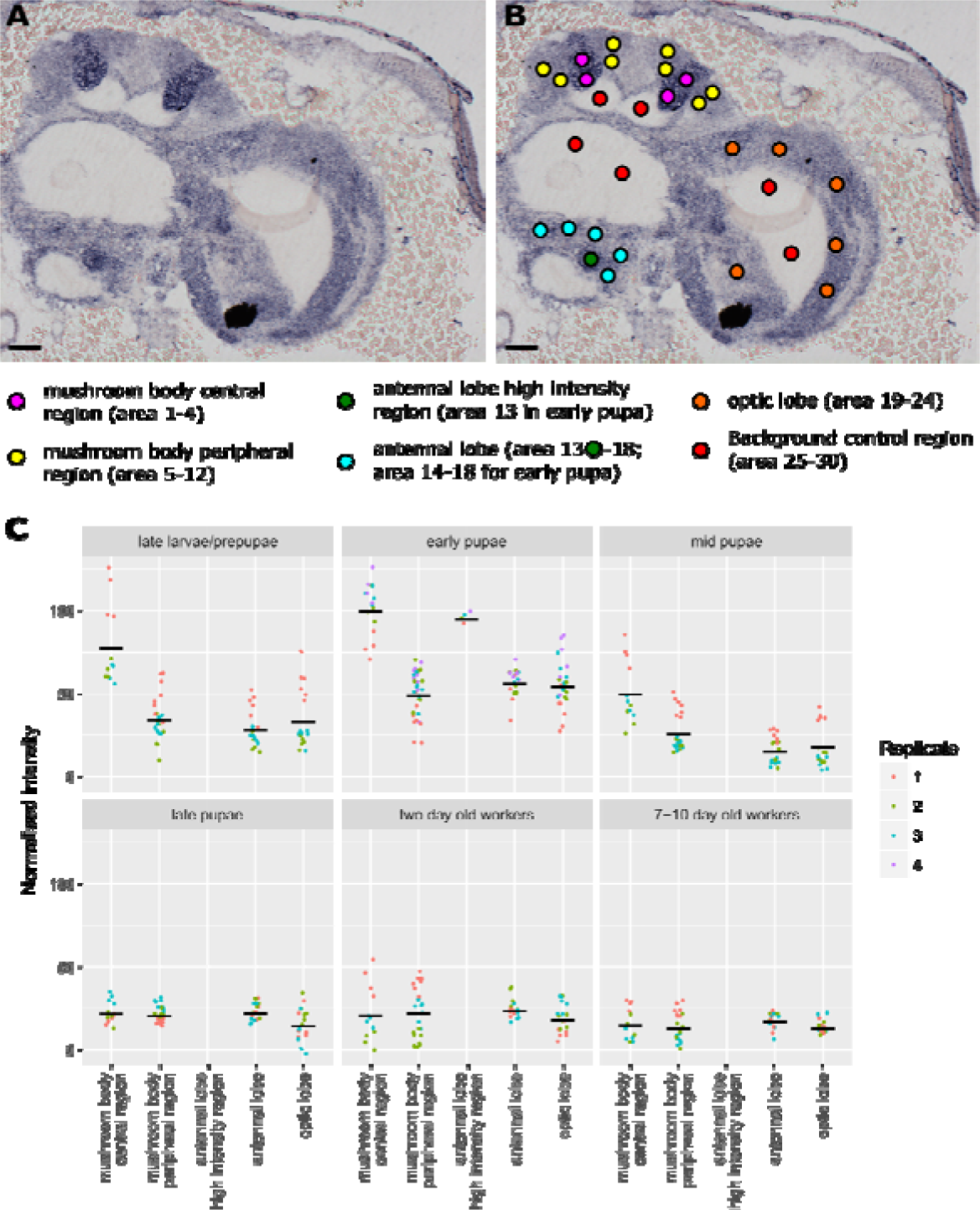
Quantification of the relative expression levels of *PRMT1* mRNA in selected areas of brains at different stages of development. A) An image example of ISH of *PRMT1* in early pupal bumblebee brain. B) Same image as in A) showing how the sub-regions to be analysed were grouped into four or five anatomical areas. C) Graph showing levels of *PRMT1* expression in brain areas (mushroom body central area, mushroom body peripheral area, antennal lobe high intensity area in early pupal stage, antennal lobe area and optic lobe area) at different stages of development. The dots indicate relative expression levels of *PRMT1*. There were three biological replicates for every stage, except for the early pupae stage where there were four biological replicates. For each stage of development six sub-regions (blue and green dots shown in B) were analysed in the antennal lobe apart from early pupal stage (five sub-regions). In early pupa, a high intensity sub-region of antennal lobe was also chosen and analysed. Different biological replicates are marked with differently coloured dots in each developmental stage. The same colour of dots in each stage were from the same individual bee. The mean of each grouped area is signified by the black bars. Scale bars: 120 μm.

**Fig. 6.**
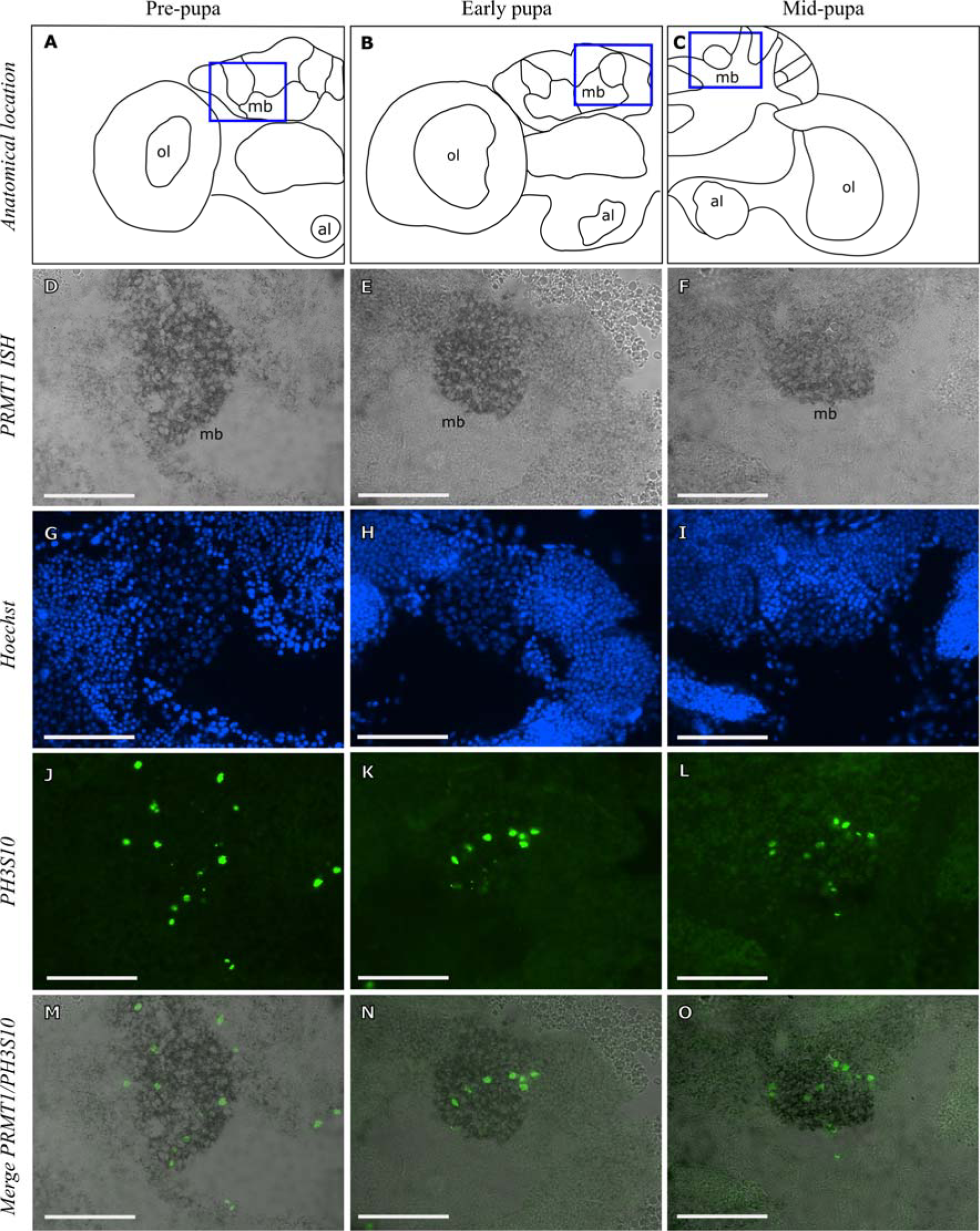
Areas of high *PRMT1* mRNA expression are enriched for dividing cells in the late larval/pre-pupal, early pupal and late pupal brain. A-C) Schematic diagrams of the late larval/pre-pupal, early pupal and late pupal brains showing the anatomical location of the mushroom bodies analysed in this study. D-F) *In situ* hybridisation of *PRMT1* in the late larval/pre-pupal, early pupal and late pupal mushroom bodies. G-I) Hoechst detection of all nuclei in the late larval/pre-pupal, early pupal and late pupal mushroom bodies. J-L) Immunohistochemical detection of mitotically active cells with anti-phospho-histone H3 Ser 10 antibody in the late larval/pre-pupal, early pupal and late pupal mushroom bodies. M-O) Merged image of *PRMT1* expression and phospho-histone H3 Ser 10 immunoreactivity in the late larval/pre-pupal, early pupal and late pupal mushroom bodies. mb: mushroom body; ol: optic lobe; al: antennal lobe. Boxes indicate the anatomic location of the mushroom bodies analysed in this study. Scale bars: 100 μm.

### Higher *PRMT1* expression correlates with mitotically active cells and lower cell density in the mushroom bodies

We quantified the relationship between higher expression of *PRMT1* and the prevalence of mitotically active cells (detected by PH3S10 immunoreactivity) within the central sub-regions of mushroom body areas in late larvae/pre-pupae, early pupae and mid-pupae. Spearman’s rank correlation coefficient was used to investigate the correlation between the expression level of *PRMT1* and the proportion of mitotically active cells (the number of PH3S10 expressing cells divided by the total number of cells as determined by the number of nuclei labelled with Hoechst) for each stage. Results presented in Fig. 7B and C showed that there was a positive correlation (Fig. 7B: *ρ* = 0.46, df = 43, p-value = 0.0015; C: *ρ* = 0.46, df = 43, p-value = 0.0015) between the proportion of mitotically active cells within a region and the level of *PRMT1* expression in that region of mushroom bodies during early and mid-pupal stages. The correlation in late larval/pre-pupal stage was weaker and not significant (Fig. 7A: *ρ* = 0.22, df = 43, p-value = 0.14), although the co-immunolabelling of *PRMT1* ISH and PH3S10 immunohistochemistry (Fig. 6M) showed that the PH3S10 positive cells were within or close to the stronger *PRMT1* ISH signal region in the central mushroom body area.

These results show that in the central mushroom body areas, at least during the early- and mid-pupae stages, high levels of *PRMT1* expression are spatially-associated with increased mitotic activity.

**Fig. 7.**
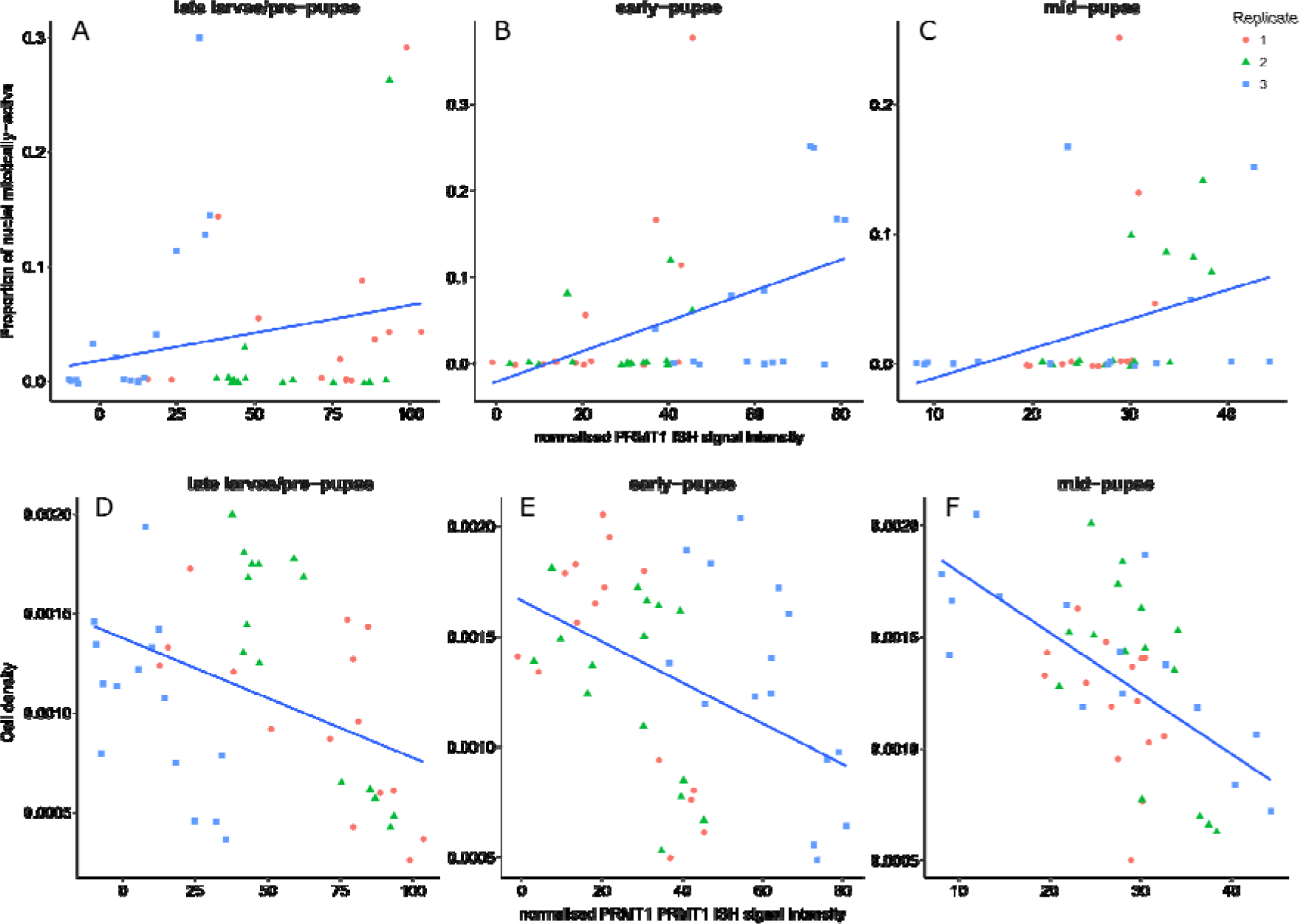
Correlation analysis of *PRMT1* expression levels and PH3S10 positive cells and cell density. A-C) Correlation between the normalised optical density of *PRMT1* ISH signals (X axis) and proportion of mitotically active cells (the number of PH3S10 expressing cells divided by the total number of cells as determined by the number of nuclei labelled with Hoechst, Y axis) in each selected region for three different stages of bees. A) 3 late larvae/pre-pupae, B) 3 early pupae and C) 3 mid-pupae. There is a moderately significant correlation between the higher proportion of dividing cells and higher expressed level of *PRMT1* for B) early pupae, *ρ* = 0.46, df = 43, p-value = 0.0015 and C) mid-pupae, *ρ* = 0.46, df = 43, p-value = 0.0015. There is no significant correlation for A) late larvae/pre-pupae, *ρ* = 0.22, df = 43, p-value > 0.14. The dots which are close to 0 value on the Y axis indicate that in these selected regions, there are no PH3S10 expressing cells. D-F) Correlation between the normalised optical density of *PRMT1* ISH signals (X axis) and cell density (total number of Hoechst stained nuclei divided by the size of the region, Y axis) in each selected region for three different stages of bees, D) 3 late larvae/pre-pupae, E) 3 early pupae and F) 3 mid-pupae. There is a stronger significant correlation between lower cell density with higher expressed level of *PRMT1* for all three developmental stages: D) late larvae/pre-pupae, *ρ* = −0.37, df = 43, p-value = 0.013; E) early pupae *ρ* =-0.4, df = 43, p-value = 0.0056; F) mid-pupae, *ρ* =-0.57, df = 43, p-value = 5.9 * 10^−5^. Each dot corresponds to the values of each selected region. The blue line is best linear fit line.

In analysing Hoechst stained images, we noticed that there were fewer nuclei in the central region of the mushroom bodies than in the outer regions of the mushroom bodies (Fig. 6G-I). In order to confirm this observation, we analysed and quantified the cell density in each region of interest. Results from Fig. 7D, E and F demonstrate that regions which show higher expression of *PRMT1* within the mushroom bodies correlated with the regions showing lower cell density (late larvae/pre-pupae/D: *ρ* = −0.37, df = 43, p-value = 0.013; early pupae/E: *ρ* = −0.4, df = 43, p-value = 0.0056; mid-pupae/F: *ρ* = −0.57, df = 43, p-value = 5.9 * 10^−5^). It is well documented that neuroblasts in the honeybee and *Drosophila* have bigger cell bodies than differentiated neurons, which may account for a lower cell density we observed in the areas of mitotic activity [1, 55, 56]. We have already shown that regions containing high *PRMT1* expression are associated with mitotically active cells. Our observation that the cell density is reduced within this part of MB further suggests that this region probably contains dividing neuroblasts [1, 13, 14].

## Discussion

In the present study, we began to investigate the roles of PRMTs in the CNS development of bumblebees by analysing the expression pattern of several *PRMT* genes in the developing brain of these insects. We found that all three *PRMTs* investigated are expressed in the developing CNS of bumblebees. The widespread expression of these genes in bumblebee brains at all developmental stages investigated suggests that they may be important throughout the life cycle of the bumblebee. Previous research into the function of PRMTs in several model organisms highlighted the multitude of processes controlled by these enzymes, such as regulation of RNA processing, DNA damage repair and signal transduction, but also their important roles in the control of neural stem cell (NSC) proliferation and homeostasis, both during development and in the adult [21, 22, 24, 45]. Our findings that the *PRMTs* are expressed throughout the life cycle of bumblebees are thus consistent with the observations from other model organisms.

Interestingly, we found that *PRMT1* was particularly highly expressed in the central mushroom body regions, which have been identified in honeybees as regions containing dividing neuroblasts/ganglion mother cells. Whilst it is not certain that areas of high *PRMT1* expression levels in the bumblebee also contain dividing neuroblasts/ganglion mother cells, such a possibility is plausible and supported by the observation that high levels of *PRMT1* expression show significant spatial correlation with the mitotic marker, PH3S10, during early and mid-pupal stages. Moreover, we observed that higher levels of *PRMT1* expression within the central mushroom body showed significant spatial correlation with regions of lower cell density. Neuroblasts in the honeybee and *Drosophila* have bigger cell bodies than differentiated neurons, which may account for the lower cell density we observed in the areas of high *PRMT1* expression characterised by the presence of mitotic activity [1, 55, 56].

These observations are intriguing given the developmental programmes taking place within the bumblebee mushroom bodies during these stages. While there is very little information about these processes in bumblebees, more is known about the developmental fates of neuroblasts in the mushroom bodies of honeybees and in *Drosophila*. Larval development in honeybees is characterised by an increase in neuroblast numbers initially, with ganglion mother cells and Kenyon cells being born towards the later larval stages until mid-late pupal stages [1]. It is noteworthy in this respect that in our study we found that in the late larval/pre-pupal stage the correlation between the level of *PRMT1* expression and the number of mitotic cells was weak, while it became stronger during the early- and mid-pupal stages. These stages correspond to two developmental events in the honeybee [1]. Firstly, during the late-larval stage the neuroblast and ganglion mother cell numbers remain high as Kenyon cells are born, suggesting a balance between proliferative and neurogenic divisions. Secondly during the early- and mid-pupal stages neuroblast and ganglion mother cell numbers decrease, whilst Kenyon cells rapidly increase in number, suggesting that neurogenic divisions predominate (See Fig. 1B) [1]. Later in development, from the mid-late pupal stage, the neuroblasts are removed by apoptosis and neurogenesis stops [1].

These observations suggest that high levels of *PRMT1* may favour neurogenic divisions of neuroblasts and ganglion mother cells to generate ganglion mother cells and Kenyon cells, respectively. This idea is further supported by the observations of mammalian cortical development, where neural stem cells that are undergoing neurogenic division to either generate intermediate basal progenitors (IBPs) or IBPs dividing to generate two neurons express high levels of the antiproliferative gene *TIS21/BTG2*, that stimulates the activity of PRMT1 [40]. Importantly, a *TIS21* orthologue is also present in bumblebees (accession number: LOC100648380; *BTG2*). Thus, the TIS21/PRMT1 axis may form part of a general mechanism controlling neurogenic divisions during development of a variety of animal species. It will be important to probe this possibility further in future studies.

In the present study we did not detect any mitotically-active cells in the late pupae and adult stages, suggesting that in the mushroom bodies of bumblebees, neurogenesis ceases during pupal development. These observations align well with previous work in honeybees, which showed that developmental neurogenesis ceases in the mushroom bodies after the mid-pupal stage, as manifested by an absence of detectable mitotically-active neuroblasts and ganglion mother cells [1, 13, 14, 57].

We cannot, however, fully exclude the possibility that adult neurogenesis may occur in bumblebees in particular situations such as brain damage as it has been observed in other insects. For example, in crickets (*Acheta domesticus*), neurogenesis in mushroom body also takes place in the adult [58]. In addition, in the medulla cortex of optic lobes of *Drosophila*, adult neurogenesis occurs as well [59]. Acute brain damage to the *Drosophila* medulla cortex triggers adult neurogenesis [59]. Investigating the possibility of such regulated adult neurogenesis in bees is an interesting goal for future work.

The *PRMT4* and *PRMT5 in situ* hybridisation results revealed a more uniform expression of both mRNAs at late larval/pre-pupal, mid- and late-pupal and adult stages. We observed a slightly stronger ISH signal in the proliferative regions of mushroom bodies in early-pupal stages for both *PRMT4* and *5* mRNAs. Interestingly, PRMT4 induces PC 12 cell proliferation by methylating the RNA binding protein HuD, and inducing degradation of anti-proliferative *p21* mRNA bound by HuD [42]. PRMT5 also maintains proliferation of PC12 cells and mouse embryonic neural stem cells during early stages of development [43-45]. PRMT5 is also required for NSC homeostasis, and its selective depletion in CNS in mice leads to CNS developmental defects and post-natal death within 14 days after birth [45]. Furthermore, *PRMT5* expression is upregulated in solid tumors, lymphoma, and leukemia [21]. Together, these observations suggest that PRMT5 may be required for maintaining cells in a proliferative state. It will be important to investigate whether PRMT5 plays a similar role in the early pupal stage in bumblebees in the future. Functional molecular studies have historically been difficult in bumblebees due to lack of genetic tools such as mutants. However new CRISPR/Cas gene editing technology has recently been applied to insects including ants and honeybees [60, 61]. The current work paves the way for such a functional study of PRMT function in bumblebees by defining the localisation and timing of *PRMT* gene expression in the brains of bumblebees.

## Data accessibility

Raw data for Fig. 5 and 7 are provided in supplementary table S1 and S2.

## Author Contributions

CG, AC and LC conceived the study. CG, AC, CJP and LC designed the study; AC, LC, ME provided technical support and advice for histology, mRNA *in situ* hybridisation, immunohistochemistry and photomicroscopy; CG performed experiments; CG and CJP analysed data; CG, AC, CJP and LC drafted the manuscript. All authors approved the manuscript.

## Competing Interests

The authors declare that they have no competing interests.

## Funding

CG was supported by a studentship from the China Scholarship Council No. 201408360067, AC was supported by the BBSRC grant BB/J006602/1. LC received support from HFSP program grant (RGP0022/2014).

## Acknowledgments.

We thank Maurice Elphick for providing access to research facilities in his laboratory and for providing feedback on the manuscript. We also thank Angelika Stollewerk, Petra Ungerer, Frederick Partridge, Thomas Butts and Weigang Cai for helpful discussions.

